# Nondimensional nucleus shape parameters reveal mechanostasis during confined migration

**DOI:** 10.64898/2026.03.24.713870

**Authors:** Ashok Ravula, Yixuan Li, Jia Wen Nicole Lee, Jia Xuan Charlotte Chua, Andrew Holle, Sreenath Balakrishnan

**Affiliations:** School of Mechanical Sciences, Indian Institute of Technology Goa, Goa, India; Mechanobiology Institute, National University of Singapore, Singapore 117411, Singapore; Department of Biomedical Engineering, College of Design and Engineering, National University of Singapore, Singapore 117583, Singapore; School of Interdisciplinary Life Sciences, Indian Institute of Technology Goa

## Abstract

Nucleus shape is a sensitive indicator of cell state, influenced by numerous biochemical and physiological factors. While prior work has cataloged how perturbations alter nucleus morphology, we address the inverse: inferring underlying molecular changes from nucleus shape alone. We previously developed a mechanical model yielding two nondimensional parameters: flatness index and scale factor, which are surrogate measures for cortical actin tension and nuclear envelope compliance respectively. In this study, we apply these parameters to investigate the dynamics in cellular mechanics during confined migration. We fabricated polydimethylsiloxane (PDMS) microchannels with widths of 3 µm (high confinement) and 10 µm (low confinement) and tracked cells migrating through them. We captured high-frequency 3D nucleus shapes via double fluorescence exclusion microscopy and custom image analysis. Fitting the model and estimating flatness index and scale factor to time-resolved shapes revealed dynamic regulation in 3 µm channels: actin tension decreased and nucleus compliance increased immediately before nucleus entry into the constriction, with rapid restoration to baseline upon exit. No such changes occurred in 10 µm channels, indicating active, confinement-dependent cytoskeletal adaptation. Immunostaining for YAP and lamin-A,C confirmed these model inferences. Our results uncover mechanostasis, active mechanical homeostasis, during confined migration and establish the combination of double fluorescence exclusion microscopy and nondimensional nucleus shape parameters as a powerful, non-invasive tool for single-cell mechanobiology studies.

## Introduction

Nucleus shape emerges from the mechanical interplay among multiple intracellular and extracellular factors such as the cytoskeletal forces, chromatin organization, the nuclear lamina and the extracellular matrix (ECM). Previous studies on nucleus shape and mechanics fall into two broad categories: (i) empirical perturbation experiments that document how targeted disruptions affect nucleus geometry [1, 2, 3, 4, 5] (ii) physics/mechanics-based models that provide theoretical frameworks for nucleus architecture [6, 7, 8]. The studies in the former category have typically characterized nucleus using purely geometric parameters such as volume, area, height, eccentricity and shape factor and cataloged changes in these parameters due to perturbations such as disrupting actomyosin contractility [5], altering ECM stiffness [1], depleting lamins [3], or modifying chromatin [4]. Illustrative examples of the latter type are a continuum-mechanics-based model that explain changes in nuclear mechanics due to modulations in cell shape and substrate stiffness [6] and a physics-based model that related the constant nucleus to cytoplasmic volume ratio observed in various cell types and organisms to the dominance of osmotic pressure in shaping the nucleus [7]. These approaches have largely remained distinct, with one emphasizing descriptive geometric quantification and the other focusing on mechanistic force-balance relations.

Previously, we had developed a description for the nucleus that is consistent with both the geometrical and mechanical points of view. Based on previous studies, we modeled nucleus as an inflated spherical balloon compressed between two rigid flat plates [8, 2]. The inflation pressure is primarily due to the osmotic pressure difference between the nucleoplasm and the cytoplasm [7, 8]. The compressive force from the top is due to the perinuclear actin cap which pushes down on the nucleus [2]. We modeled this configuration using mechanics of membranes by assuming the nuclear envelope as a hyperelastic membrane [9, 10]. The nuclear envelope here refers to a combination of the nuclear double membrane and the underlying lamina. By using this model, we derived two nondimensional parameters that can characterize nucleus shape, namely, scale factor and flatness index [11]. Independent geometric variability analysis of around a thousand nuclei from several cell lines showed that the two dominant variability modes are scaling and flattening, which together account for 85% of the variability [12]. We further showed that these variability modes align directly with the mechanically derived parameters, thereby uniting both the geometrical and mechanical points of view. Further these nondimensional quantities enable inverse inference: extracting information on underlying mechanical properties (actin tension and nuclear compliance) from observed shape changes.

The dominant force on the nucleus envelope is the osmotic forces that tends to inflate (scale) the nucleus. The lamin meshwork, which confers stiffness to the nucleus envelope, resists this scaling and hence the scale factor correlates with nucleus compliance. Therefore, scale factor inversely correlates with lamin-A,C, which is the major load-bearing member of the nucleus envelope. The tension in cortical actin compresses the nucleus from the top through the perinuclear actin cap leading to nucleus flattening [2]. Hence, the flatness index correlates with actin tension. We have experimentally demonstrated these relations on several cell lines through multiple perturbations such as actin and microtubule depolymerizing drugs [11], novel Myosin Light Chain Kinase inhibitors [13], Hepatitis C Virus [10] and modifying the elasticity of the cell substrate [11]. Here, we apply this framework to investigate dynamic mechanoregulation during cell migration through physical confinements.

Cell migration is an important process during various physiological phenomena such as embryonic development [14, 15], wound healing [16], immune response [17] and tissue regeneration [18]. In the body, cells need to migrate through the interstitial space (size 0.1 - 30 µm), which is smaller than or comparable to the size of migrating cells [19]. To penetrate such confinements the cells need to remodel their shape and modify their mechanical properties. Such modulation of cellular properties is accomplished through biochemical and biomechanical feedback mechanisms [20, 21]. Confined cell migration has also been shown to directly influence cell fate decisions [22], pathological conditions [23], fibroblast activation [24], and be involved in cellular mechanical memory [25]. Several previous studies have focused on identifying the changes in cytoskeletal components accompanying confined migration [26, 27, 28]. An important question is how the nucleus, which is the stiffest organelle, traverses the confinement. Further, the nucleus is known to soften during confined migration by down-regulating lamin-A,C [29, 30, 31]. Along with the reduction in nucleus stiffness, the cell stiffness is also known to decrease during confined migration [32]. Since cell stiffness is predominantly due to cytoskeletal contractility, the reduction in cell stiffness suggests reduction in actin tension [33, 34]. However, there are several unanswered questions such as (i) at which stage during the confined migration are these biomechanical properties modulated, (ii) do these properties revert to their native state after the cells exit the confinement and (iii) do these dynamics depend on the severity of confinement?

To investigate the aforementioned questions, we have studied the morphology of nuclei when the cells migrate through 3 and 10 µm wide channels (Fig. 1). We fit the nucleus shapes to our mechanical model and calculate flatness index and scale factor to infer the dynamics of actin tension and nucleus compliance. Our results show that cells decrease the actin tension and increase nucleus compliance, in a confinement-dependent manner, immediately before the nucleus enters the channel and further these changes recover once the nucleus exits the channel. Such temporary perturbation of these biomechanical parameters due a physical challenge and recovery post challenge suggests mechanostasis, a portmanteau of mechanical and homeostasis.

**Figure 1.**
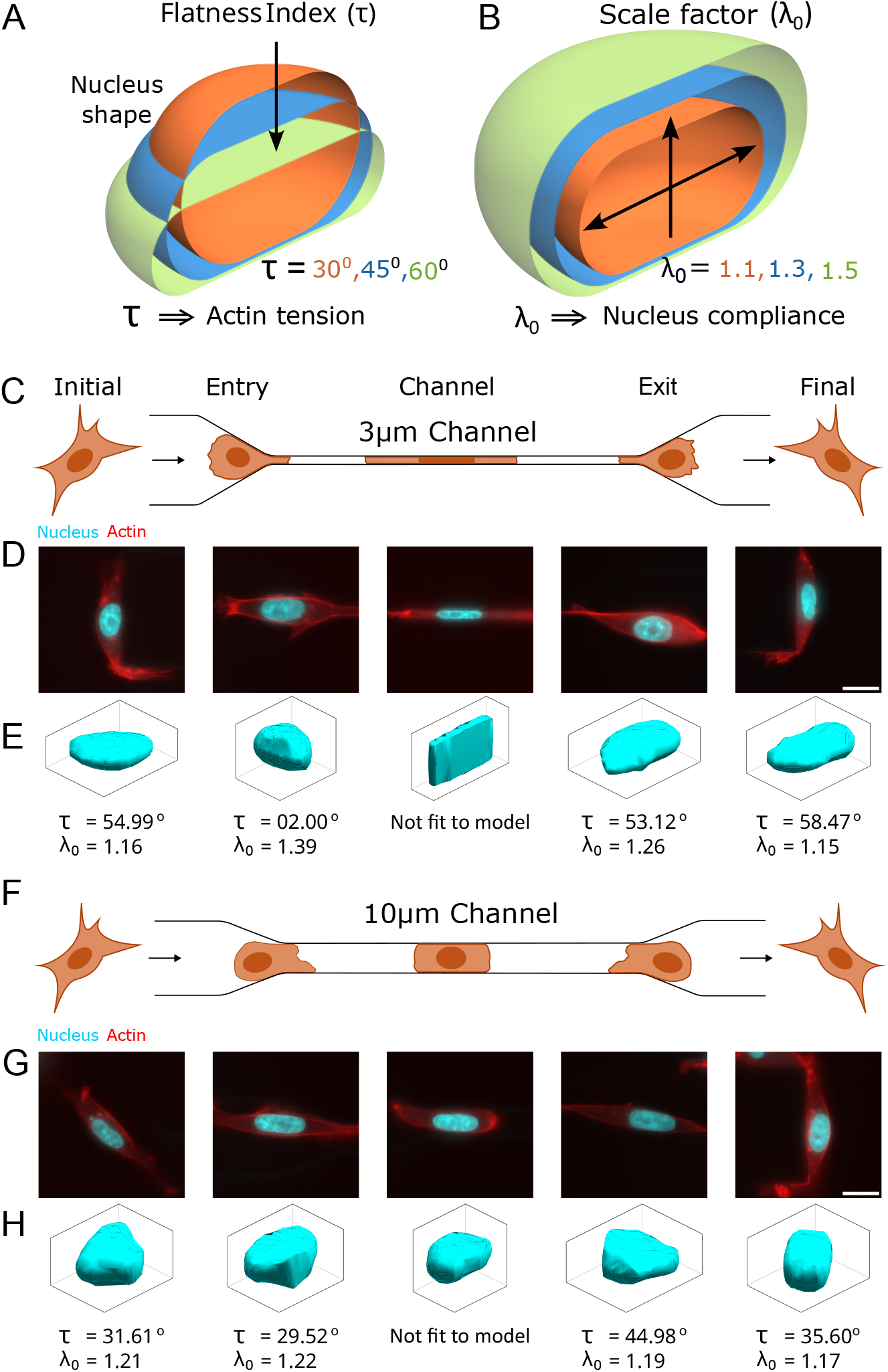
Nucleus shape remodelling during confined migration. Geometric interpretation of nucleus nondimensional parameters (A) Flatness index and (B) Scale factor. (C) Schematic of cell migration through 3µm channel. Five positions are shown: (i) initial - cell has not entered the channel, (ii) entry - part of the cell has entered the channel, but the nucleus is completely outside the channel, (iii) channel - cell and nucleus is completely inside the channel, (iv) exit - nucleus has completely exited the channel, but part of the cell is remaining inside the channel and (v) final - the cell has completely exited the channel. (D) Representative fluorescence images of cells at the aforementioned positions (E) Representative 3D nucleus shapes obtained using double-fluorescence exclusion microscopy with flatness index and scale factor determined by fitting the nucleus shape model. (F) Schematic, (G) fluorescence images and (H) nucleus shapes for migration through 10 µm channel. Scale bar is 15 µm.

## Materials and Methods

### Cell culture

MDA-MB-231 cell line was obtained from ATCC and routinely tested for Mycoplasma contamination. Cells were maintained in a humidified incubator at 37 °C with 5% CO_2_ and were passaged once cultures reached approximately 70–80% confluence. Cells were cultured using Dulbecco’s Modified Eagle Medium (DMEM; GIBCO), supplemented with 10% Fetal Bovine Serum (FBS; GIBCO), and 1% Penicillin-Streptomycin (GIBCO).

### Microchannel Chip Fabrication and Confined Migration Experiment

Silicon wafers containing custom microchannel arrays measuring 150 µm in length, 10 µm in height, and either 3 µm or 10 µm in width were fabricated by the Microfabrication Core at the Mechanobiology Institute (MBI). The silicon masters were generated using a two-step photolithography process. Wafers were first dehydrated at 180 °C for 15 minutes and then spin-coated with a 10 µm layer of SU-8 3010 photoresist (2900 rpm, 30 s). The coated wafers were soft-baked (65 °C for 1 min, 95 °C for 5 min), exposed to 110 mJ/cm^2^ UV light through a sodalime optical mask (SUSS Microtec MJB4, 365 nm filter), and post-baked (65 °C for 2 min, 95 °C for 5 min). Following development using SU-8 developer, the wafers were rinsed with isopropyl alcohol, dried with nitrogen gas, and hard-baked at 130 °C for 5 minutes with cooldown steps at 95 °C and 65°C. A second SU-8 3050 layer (100 µm thickness) was subsequently spin-coated (1200 rpm, 30 s), soft-baked (65 °C for 10 min, 95 °C for 20 min, 65 °C for 10 min), exposed to 150 mJ/cm^2^ UV light, and post-baked (65 °C for 2 min, 95 °C for 5 min, 65 °C for 2 min). After development, the wafers were rinsed, dried, and hard-baked. All SU-8 photoresists and developer were obtained from Kayaku Advanced Materials (MA, USA). Sylgard 184 polydimethylsiloxane (PDMS; Dow-Corning) was prepared by mixing the polymer base with a curing agent at a 10:1 ratio, followed by degassing under vacuum for 5 minutes to remove air bubbles. The PDMS mixture was then poured over the silicon wafer mold and cured at 80 °C for 2 hours before the negative microchannel structures were excised from the wafer. The patterned PDMS and glass coverslips were plasma treated and bonded to form microchannel devices. These chips were then coated overnight at 37 °C with 0.3 mg/mL rat-tail collagen-1 (GIBCO) dissolved in 0.1% acetic acid. Cells were seeded into the entrance reservoir of the microchannel devices at a density of 1 × 10^6^ cells/mL, and allowed to migrate through the channels for 24 hours.

### Immunofluorescence and Microscopy

Cells were fixed with 3.7% paraformaldehyde (PFA) at room temperature for 45 minutes, followed by permeabilization and blocking with 0.1% Triton X-100 and 1% bovine serum albumin (BSA) in PBS for 1 hour at room temperature. For immunostaining, both primary and secondary antibodies were diluted in a 1% BSA solution containing 0.1% Tween-20 in PBS (PBST). Cells were incubated overnight at 4 °C with the primary antibodies against lamin A/C (Abcam, 1:250, ab40567) and YAP (Santa Cruz Biotechnology, 1:100, sc-101199). After three washes with 0.1% PBST, cells were incubated for 4 hours at room temperature with the secondary antibody solution (Invitrogen, Donkey anti-Rabbit IgG (H+L) Alexa Fluor™ Plus 568 or Donkey anti-Mouse IgG (H+L) Alexa Fluor™ Plus 568, 1:1000) and Hoechst 33342 (Invitrogen, 1:5000, H1399). Fluorescence microscopy was performed using the ZEISS Celldiscoverer 7 microscope with a Plan-APOCHROMAT 20×/0.7 NA autocorrective objective.

### Mechanical model for nucleus shape

Based on previous studies [8], we modeled the nucleus as an inflated spherical membrane compressed between two rigid plates [10]. The nuclear envelope was modeled as an incompressible Mooney-Rivlin material, which is spherical in the unloaded state. By analyzing this mechanical configuration using the mechanics of membranes [9], we showed that the nucleus shape can be characterized using two non-dimensional parameters - (i) flatness index, *τ*, which represents the flattening and (ii) scale factor, *λ*_0_, which represents isometric scaling [11]. These parameters were shown to form a canonical basis through variability analysis of nucleus shapes from multiple cell lines [12]. Through various experiments involving depolymerization of actin filaments and microtubules, inhibition of Myosin Light Chain Kinase, growing cells on substrates of varying stiffness and Hepatitis C Virus infection of liver cells we showed that flatness index corresponds to actin tension and scale factor corresponds to nucleus compliance [10, 11, 13]. Here, we fit this model to the nucleus shapes during confined migration, estimate flatness index (*τ*) and scale factor (*λ*_0_) at each time point and thereby infer the changes in actin tension and nucleus compliance during confined migration.

### Statistical analysis

Statistical analysis was performed using unpaired Student’s t-test (* - p value *<*0.05, ** - p value *<*0.01, and *** - p value *<*0.001).

## Results

To investigate the dynamics of actin tension and nucleus compliance during cell migration, the cell is allowed to traverse through the microchannels of widths 3 µm and 10 µm (Fig. 1D and G). During this migration, the nucleus shapes were measured using the double fluorescence exclusion microscopy method [35]. From these images, we reconstructed the 3D shapes of the nucleus and further calculated the nucleus shape parameters, flatness index (*τ*) and scale factor (*λ*_0_), to infer actin tension and nucleus compliance respectively (Fig. 1). A representative plot showing the fluctuations in these parameters during migration is shown in Fig. 2. When the nucleus is inside the channel, it experiences lateral forces from the channel walls. Since these forces are not considered in the nucleus shape model, we cannot fit the model when the nucleus is inside the channel. From the rest of the cell positions, we chose four positions for comparison—(i) Initial: cell has not entered the channel, (ii) entry: part of the cell has entered the channel, but the nucleus is completely outside the channel, (iii) exit: nucleus has completely exited the channel, but part of the cell is remaining inside the channel and (iv) final: the cell has completely exited the channel (Fig. 1).

**Figure 2.**
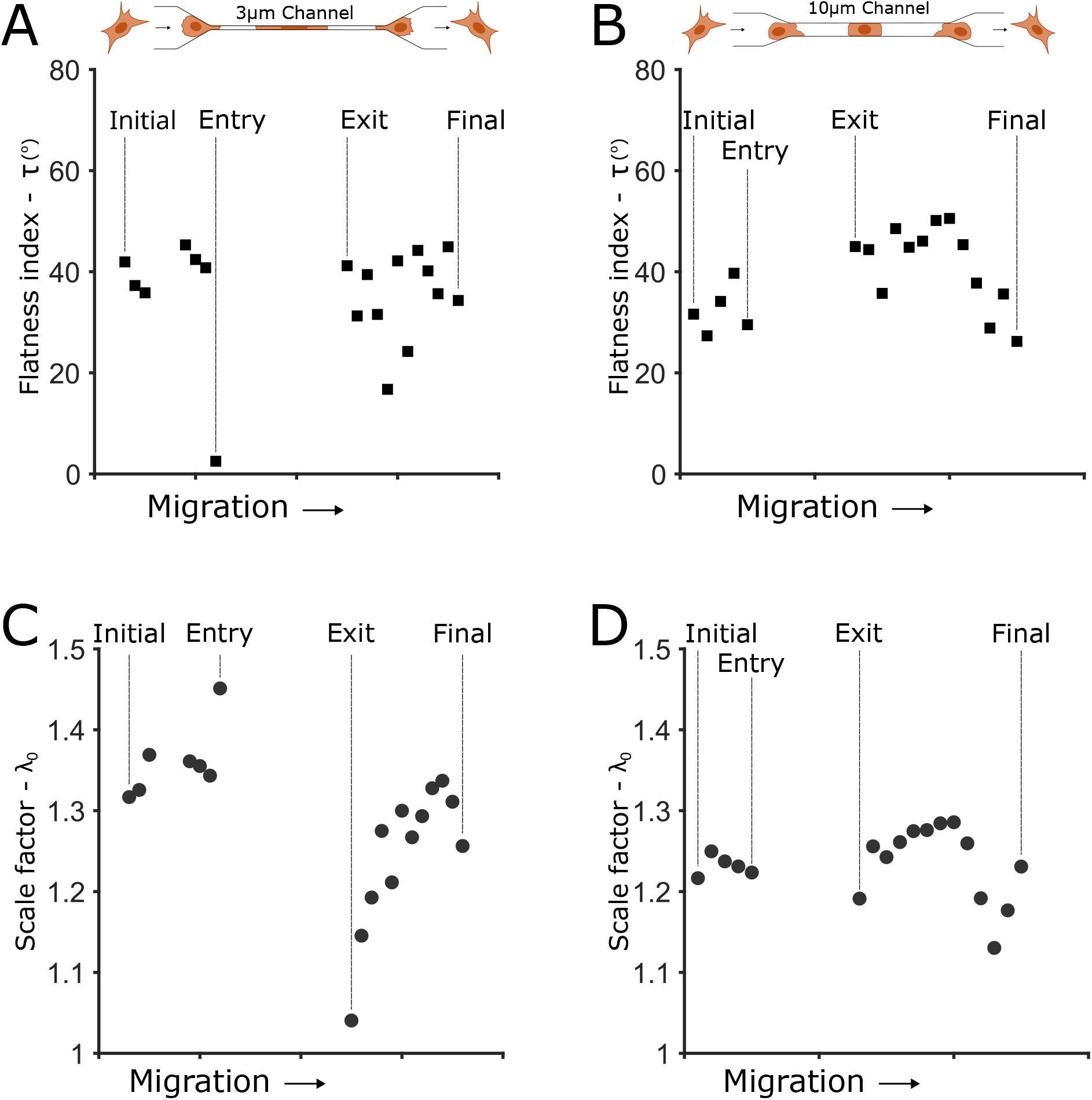
Representative plot of variation in flatness index (*τ*) and scale factor (*λ*_0_) during confined migration through 3 µm(A,C) and 10 µm(B,D) channels.

### Dynamics of actin tension during confined migration

Flatness index at initial, entry, exit and final for 3 µm and 10 µm channels are shown in Figure 3 A and B respectively. For 3 µm channel, flatness index decreased significantly at the channel entry, recovered partially at exit and further recovered completely towards the end (Fig. 3A). In case of 10 µm channel, while there is a similar trend, reduction at entry, partial recovery at exit and complete recovery at the end, the changes are not significant (Fig. 3B). Hence, our model predicts confinement-dependent, dynamic modulation of actin tension during confined migration.

**Figure 3.**
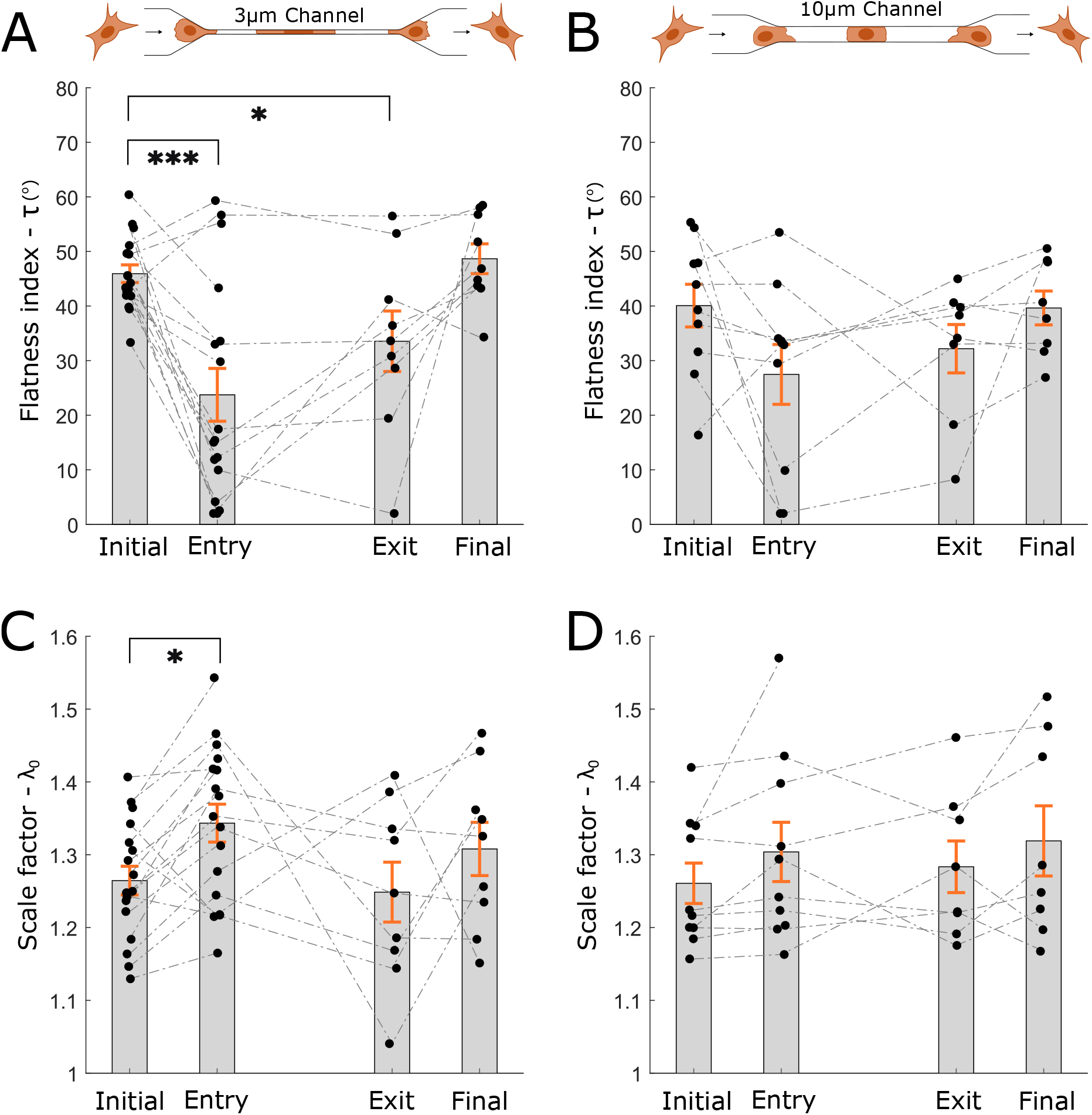
Variation of flatness index and scale factor during confined migration. Flatness index (*τ*) and scale factor (*λ*_0_) at initial, entry, exit and final for 3 µm (A and C) and 10 µm (B and D) channels. The dotted lines connect the same cell at different positions. Statistical analysis was done using unpaired Student’s t-test (* - p*<*0.05, ** - p*<*0.01 and *** - p*<*0.001).

To experimentally validate this prediction, we stain for Yes-Associated Protein (YAP) on migrating cells (Fig. 4). Nuclear localization of YAP is known to indicate high actin tension whereas translocation of YAP to the cytoplasm indicates low actin tension [36, 37, 38]. Hence, we quantified actin tension by calculating the nuclear-to-cytoplasmic ratio (N/C) of YAP intensity. N/C was calculated at the four positions defined earlier: (i) Initial, (ii) entry, (iii) exit, and (iv) final. In case of 3 µm channels, N/C decreased significantly at entry, partially recovered at exit and fully recovered when the cell completely exited the channel (Fig. 4 A and C). In contrast, for the 10 µm channels N/C did not differ among initial, entry, exit and final (Fig. 4 B and D). These immunofluorescence results confirm the model’s prediction that the cells transiently downregulate and subsequently restore actin tension during confined migration in a confinement-dependent manner.

**Figure 4.**
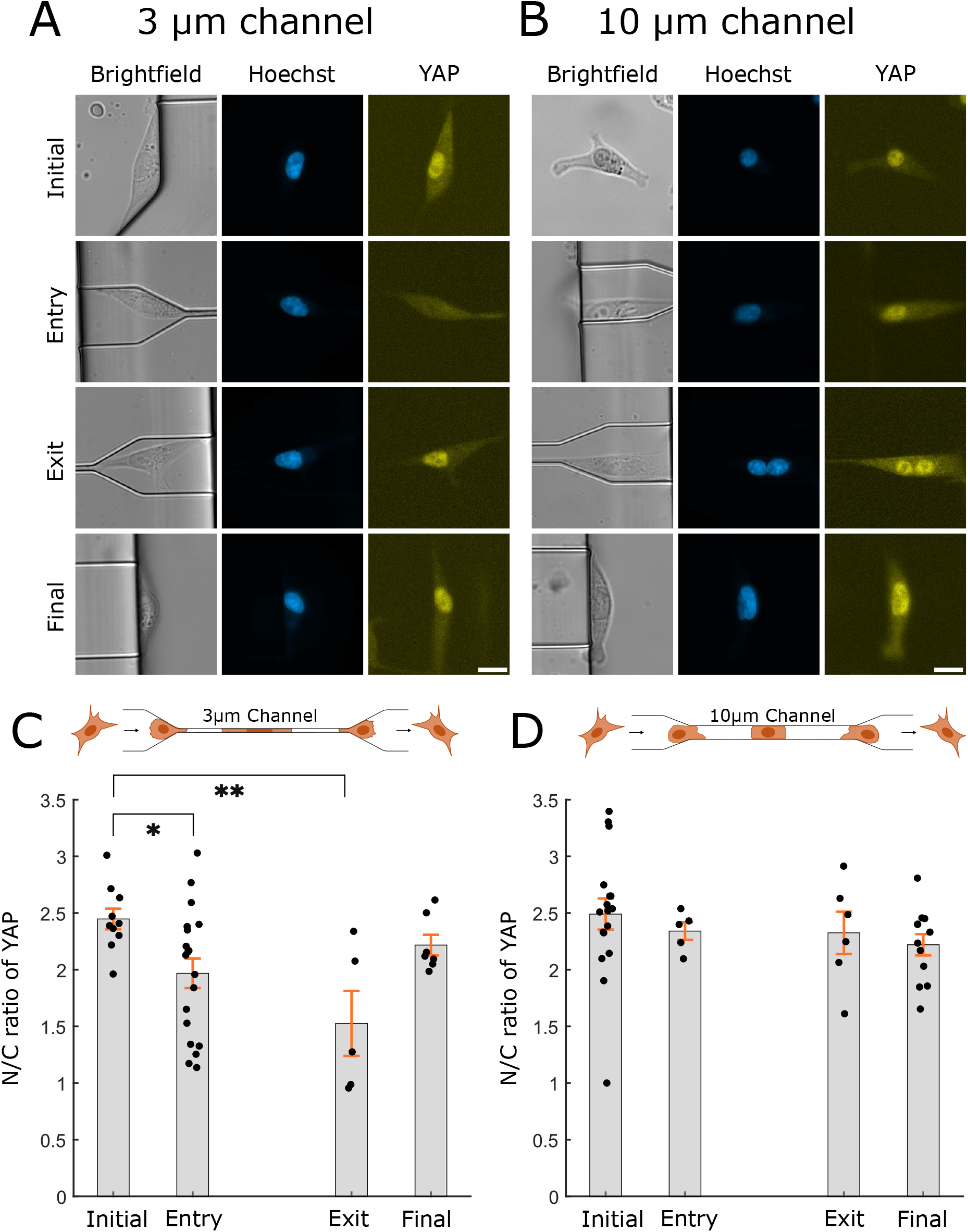
YAP translocation during confined migration. Nuclear localization of YAP indicates high actin tension, whereas cytoplasmic localization indicates low actin tension. Hence, nucleus to cytoplasm ratio of YAP (N/C) is an indirect measure of actin tension [36, 37, 38, 39]. Brightfield and fluorescence images of cells migrating in (A) 3 µm and (B) 10 µm channels. The nucleus is shown in blue and YAP in yellow. N/C at initial, entry, channel, exit and final for (C) 3µm and (D) 10µm channels. The scale bar is 15 µm. Statistical analysis was done using unpaired Student’s t-test (* - p*<*0.05, ** - p*<*0.01 and *** - p*<*0.001).

The first part of our observations, the reduction in actin tension during confinement, is supported by several previous studies. By using Atomic Force Microscopy (AFM), Rianna et al. showed that cell stiffness decreases with increasing confinement [39]. Since actin tension is the major contributing factor for the cell stiffness, this reduction in cell stiffness suggests reduction in actin tension [33, 34]. Aberrations in the actin cytoskeleton architecture under confinement, suggesting loss of tension, was also observed using immunofluorescence imaging [40, 41, 42]. Traction forces exerted by cells were shown to be significantly lower in higher confinement compared to lower confinement. For this experiment, a specialized microfluidic confinement channel with micropillars was used [43]. Since traction forces are due to actin tension, this observation concurs with our results. Altered actin network and cytoplasmic localization of YAP in the cells in narrower channels further confirmed the reduced contractility [39]. Cell softening was also shown during trans-endothelial migration using Brillouin confocal microscopy [44]. All the studies mentioned above independently show that actin tension is reduced during confined migration. The second part of our observations, recovery of actin tension after exiting from confinement, is a new result from our study.

### Dynamics of nucleus compliance during confined migration

Similar to flatness index, we compare scale factor at initial, entry, exit, and final positions. In the case of 3 µm channel, scale factor significantly increased at channel entry (Fig. 3C). This enhancement recovered completely at the channel exit, which further did not change towards the end (Fig. 3C). In the case of 10 µm channel, there is no significant change in scale factor during the migration (Fig. 3D). Since scale factor correspond to nucleus compliance, our results show that cells actively modulate the stiffness of their nuclear envelope during confined migration. When the cell anterior encounters the channel (at entry), it increases nucleus compliance, possibly by downregulating lamin-A,C, to ease the traversal through the channel. Once the cell exits the channel, it recovers to its earlier nucleus compliance by upregulating lamin-A,C. By comparing between the channels, we observe that this modulation depends on the degree of confinement. For higher confinement (lower channel width - 3 µm), the cell increases nucleus compliance whereas for lower confinement (higher channel width - 10 µm), there is no enhancement. This suggests that when the leading edge at the cell anterior confronts the channel, it is able to assess the degree of confinement and appropriately regulate nucleus compliance to aid migration.

To verify these predictions experimentally, we stained for lamin-A,C (Fig. 5). Similar to the previous analysis of actin tension (Fig. 4), we compared lamin-A,C intensity at four positions—(i) initial, (ii) entry, (iii) exit and (iv) final. In case of 3 µm channel, lamin-A,C decreased significantly at entry, recovered at exit and did not further change towards the end (Fig. 5C). On the other hand, in the 10 µm channel, even though there was some decrease in lamin-A,C at entry, it was lower than the decrease in case of 3 µm channel (Fig. 5D). Since, lamin-A,C confers stiffness to the nucleus envelope, inverse of nucleus compliance, these experimental results confirm our model-based inferences (Fig. 3C and D).

**Figure 5.**
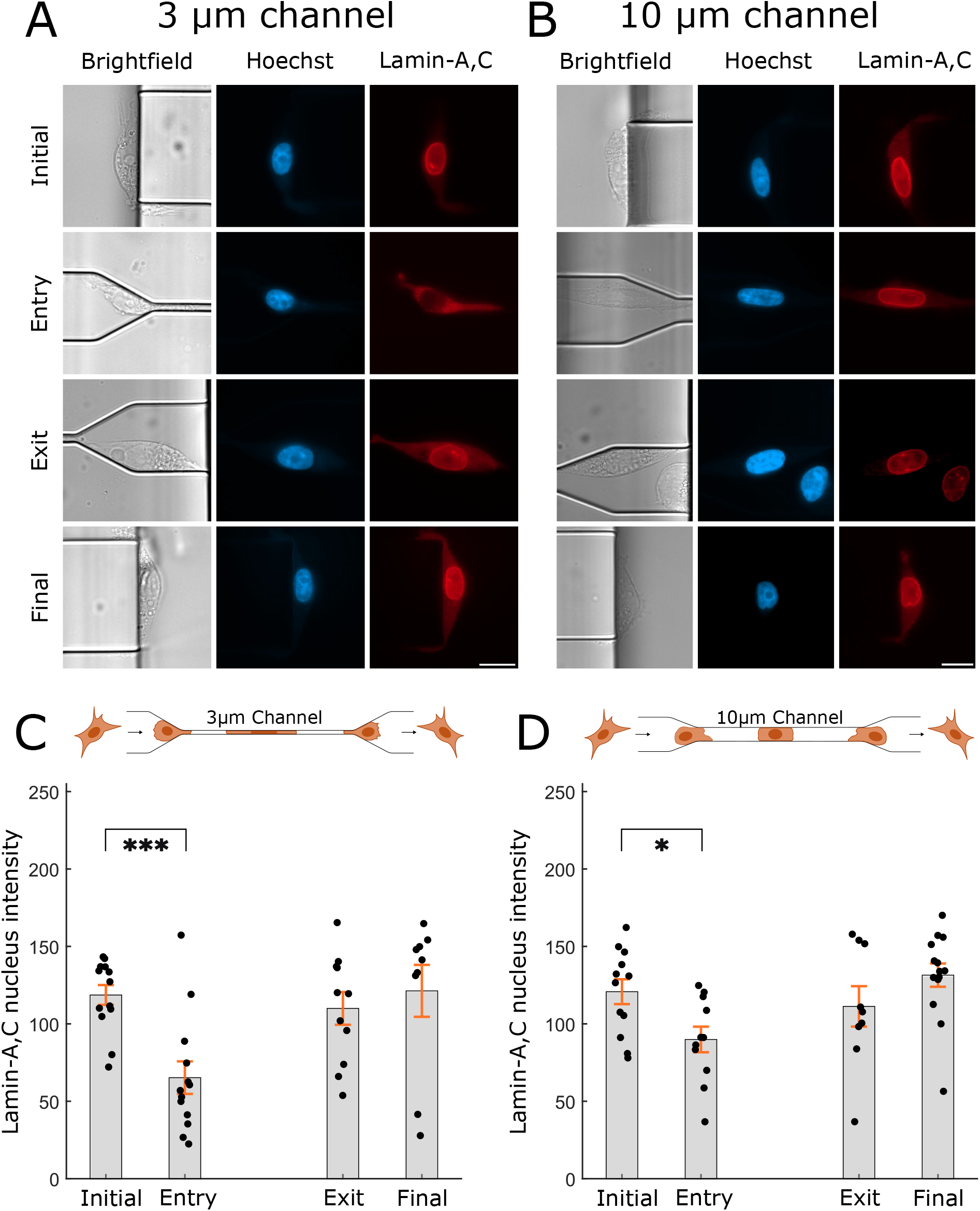
Modulation of lamin-A,C during confined migration. Reduced fluorescence of lamin-A,C indicates enhanced nuclear compliance (reduced nuclear stiffness). Brightfield and fluorescence images of cells migrating in (A) 3 µm and (B) 10 µm channels. The nucleus is shown in blue and lamin-A, C in red. Lamin-A,C intensity at initial, entry, channel, exit, and final positions for (C) 3 µm and (D) 10 µm channels. The scale bar is 15 µm. Statistical analysis was done using unpaired Student’s t-test (* - p*<*0.05, ** - p*<*0.01 and *** - p*<*0.001).

Since the nucleus is the stiffest cell organelle, it is the primary barrier to confined migration. Hence, it is prudent from the cell’s point of view to enhance nucleus compliance to facilitate migration. Increasing nucleus compliance, by downregulating lamin-A,C, was shown to enhance migration speeds through constrictions and vice versa [45]. In [46], the authors analyzed the proteome of cells inside 3 and 15 µm wide channels and found that the cells in the 3 µm channel had significantly lower levels of lamin-A,C in comparison to those in the 15 µm channel. Another independent study showed that perinuclear lamin meshwork was fragmented in 2 µm wide channels compared 8 µm wide channels [47]. Since lamins are the primary contributor to nucleus stiffness, these results suggest nucleus softening during confined migration. Furthermore, such nucleus softening was independently demonstrated using Brillouin confocal microscopy [44]. While these studies supports our first observation of increasing nucleus compliance at entry, our second observation of restoring nucleus compliance upon exit adds a new dimension to the inter-relationship between lamin-A,C and confined migration.

## Discussion

By using our mechanical model [11], we had previously predicted alterations in nuclear mechanics due to Hepatitis C Virus [10] and screened small molecule inhibitors for Myosin Light Chain Kinase [13]. Here, we applied this model to understand mechanical adaptation of single cells to physical confinement. By fitting the model to nucleus shape data, we obtained two nondimensional parameters, flatness index (*τ*) and scale factor (*λ*_0_), at several positions during confined migration. Since flatness index corresponds to actin tension and scale factor corresponds to nucleus compliance, we could monitor the variation of these important biophysical properties during confined migration. We observed that actin tension decreases and nucleus compliance increases when the cell encounters the channel and subsequently recovers after the nucleus has exited the channel, demonstrating mechanostasis during confined migration (Fig. 6). Furthermore, this dynamic modulation is dependent on the severity of confinement. In the case of channels with higher width (lower confinement), the degree of modulation is lower (Fig. 6). Immunofluorescence imaging of YAP and lamin-A,C confirmed the inferences from the model (Fig. 6). The excellent agreement between the model and experiments demonstrate that cells transiently regulate actin tension and nucleus compliance while traversing through confinements.

**Figure 6.**
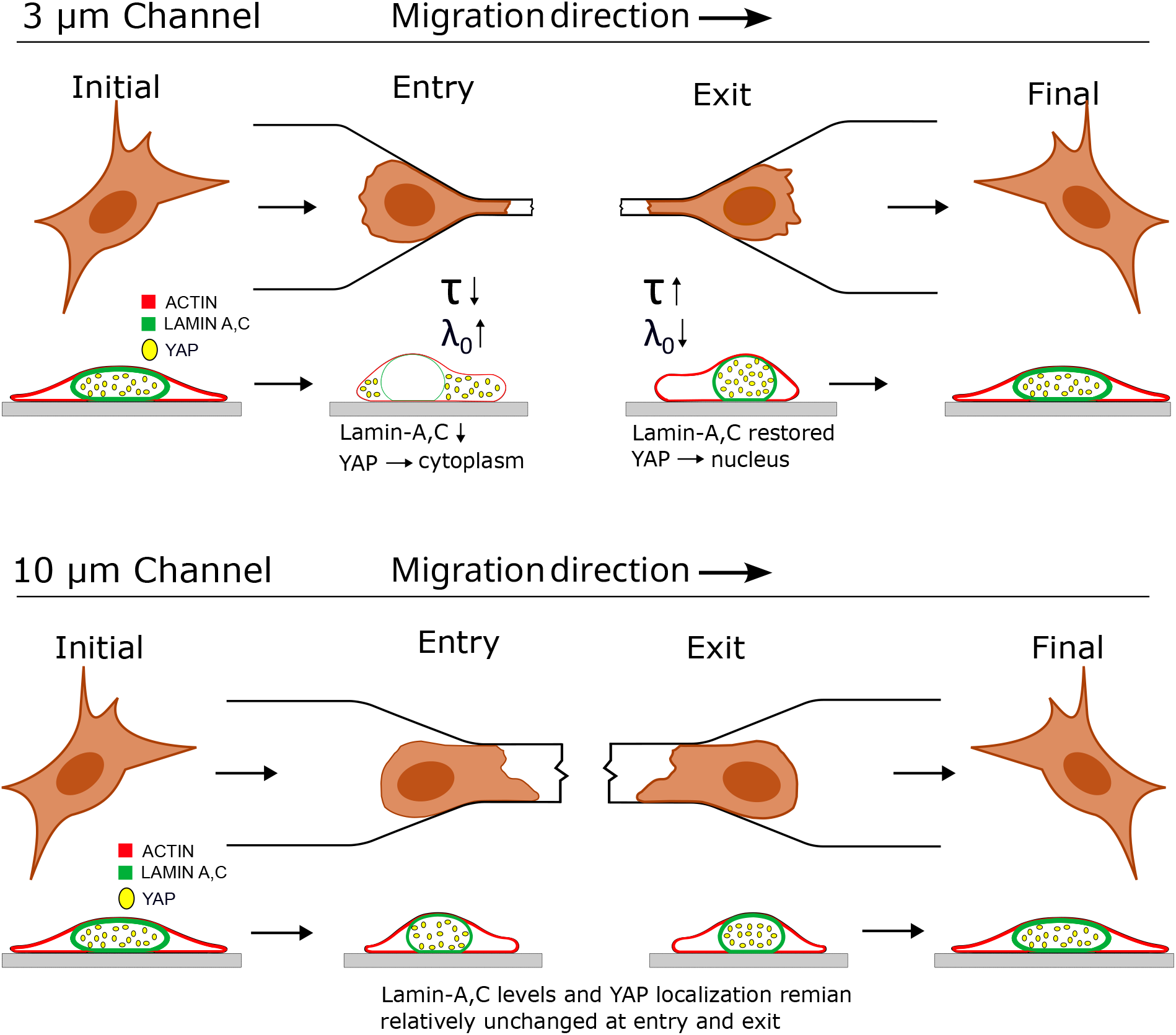
Schematic summarizing the variations in actin tension and nucleus compliance during confined migration. By analyzing the alterations in nucleus shape during confined migration using our nucleus shape model, we predicted the following for 3 µm channel: (i) actin tension reduced at entry and recovered at exit and (ii) nucleus compliance increased at entry and recovered at exit. In case of 10 µ channel, the modulations were of much lower degree. These predictions were confirmed by YAP nuclear localization (reflecting actin tension changes) and lamin-A,C staining (reflecting nucleus compliance changes).

In addition to the modulations of actin tension and nucleus compliance before the nucleus enters the channel, which is supported by the many independent experimental observations, our results show that these perturbations recover after the nucleus exits the channel. Furthermore, these modulations are proportional to the degree of confinement and are triggered when the leading edge of the cell has entered the channel and the nucleus is completely outside the channel. This suggests that the leading edge of the cell is detecting the degree of confinement and appropriately modifying the cell properties to facilitate migration through the narrow channel. Conversely, when the nucleus has exited the channel, the cell detects the absence of confinement and recovers the actin tension and nucleus compliance. Such an ability to recover from perturbations to its initial mechanical state suggests mechanical homeostasis mechanisms, which we have termed as mechanostasis, within the cellular machinery.

Mechanostasis has been described in a range of biological contexts such as apoptosis and disease [48], to describe the ability of cells to maintain or restore a preferred mechanical state despite external perturbations, through coordinated regulation of cytoskeletal tension, nuclear structure, and mechanotransduction pathways. In this context, the behavior observed here during confined migration represents a functionally important form of mechanostasis. Cells navigating confining environments must temporarily reduce actomyosin tension and increase nuclear compliance to permit nuclear passage, but these changes cannot be sustained without functional consequences. Cells such as immune cells and fibroblasts routinely migrate through dense, confining tissues, and repeated transit through confinement would be expected to impose significant mechanical stress on both the cytoskeleton and nucleus. Without the ability to restore baseline mechanical properties after each migration event, cells would accumulate damage as a result of nuclear envelope rupture, DNA damage, or persistent alterations to their chromatin organization, ultimately compromising cellular function. Mechanostasis therefore describes the processes that cells undergo to balance the need for deformability during migration and the requirement to preserve nuclear integrity and mechanical stability over time.

While several previous studies have shown the modulation of actin cytoskeleton and lamin meshwork prior to cells entering confinement, they have employed sophisticated experimental methods such as AFM, proteomics and Brillouin confocal microscopy to arrive at these conclusions. In comparison, we have used only the nucleus shape and its associated two nondimensional parameters, flatness index and scale factor, to arrive at similar inferences and further uncover mechanostasis upon channel exit. This demonstrates the power of combining double fluorescence exclusion imaging and the nucleus shape model for convenient, quick and deep inferences for morphology-based single-cell studies.

Nuclear morphology have long served as established diagnostic and prognostic indicators in pathology. The Papanicolaou (Pap) smear exemplifies this principle, where deviations in nuclear size, shape, contour irregularity, and chromatin distribution enable detection of cervical dysplasia and carcinoma [49]. Aberrant nuclear features such as irregular contours, blebbing or increased nuclear-to-cytoplasmic ratio are hallmarks of malignancy across numerous solid tumors (including breast, lung, prostate, ovarian, liver, and thyroid carcinomas) and are integral to histopathological grading [50, 51]. In laminopathies, including Hutchinson–Gilford progeria syndrome, Emery–Dreifuss muscular dystrophy, dilated cardiomyopathy, and familial partial lipodystrophy, mutations in nuclear lamina proteins (notably lamin A/C) result in nuclear envelope instability, which manifests as morphological defects that impair mechanotransduction and contribute to genomic instability [52, 53, 54]. Although these morphology-based assessments have proven clinically valuable, they are typically context-specific, applied within defined disease states or cell lineages and rely on qualitative descriptors tailored to particular pathologies [50, 54]. In contrast, the present framework provides a generalized, biophysically grounded approach that quantitatively links nuclear morphology to cell mechanical properties, namely, perinuclear actin cortical tension and nuclear envelope compliance, across diverse adherent cell types. By deriving two nondimensional parameters (flatness index and scale factor) from a reduced-order hyperelastic membrane model, we enable non-invasive inference of these properties directly from shape measurements, independent of disease or cell-type specificity. This broad applicability is supported by consistent performance of the model across multiple cell lines (Huh7, MCF7, MDA-MB-231, NIH-3T3, HeLa) under varied perturbations, including hepatitis C virus infection [10], cytoskeletal inhibitors [11, 13], and substrate stiffness modulation [11]. In the current work, we extend its utility to dynamic contexts by demonstrating its ability to capture confinement-dependent mechanoregulation during migration, further underscoring its potential as a versatile tool for interrogating cellular mechanics in physiological and pathological settings.

We previously demonstrated that variability analysis can reveal the relative importance of different contributors to nucleus morphology [12]. The first two principal modes of nucleus shape variation, scaling and flattening, together account for approximately 85% of the total observed variability in nucleus shapes. This dominance allowed us to construct a computationally efficient, reduced-order model that incorporates only these two modes yet faithfully approximates experimentally measured nucleus morphologies. Variability analysis further enables systematic identification of subsequent important modes, guiding meaningful model extensions. The third-ranked mode, eccentricity (accounting for *≈* 4% of shape variability), can be incorporated by relaxing the assumption of axisymmetry in our current framework. The mechanical component of our approach employs a reduced-order membrane mechanics model, which occupies a valuable intermediate position between overly simplistic lumped-parameter models and computationally demanding full continuum models. This formulation yields nondimensional parameters that are physically interpretable and strongly predictive. Taken together, our results indicate that combining variability analysis with reduced-order physical modeling offers a powerful, viable strategy for developing mechanistic inference models in biology.

